## Supplementary Information for "Modality-specific tracking of attention and sensory statistics in the human electrophysiological spectral exponent"

### Supplemental Material

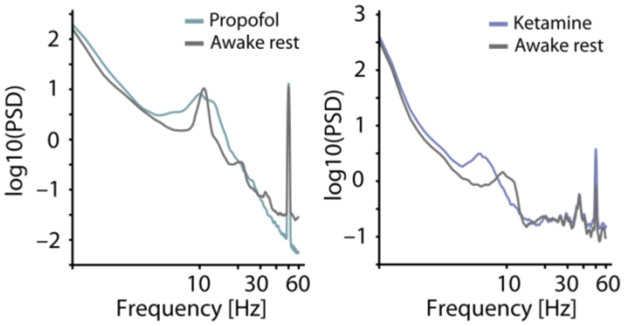


**Figure 1 supplement 1:** Raw EEG power spectra during awake rest, ketamine, and propofol anaesthesia. Non-normalized EEG spectra averaged across 5 subjects and 5 central electrodes (inset) displaying a contrast between rest and propofol (left) and ketamine anaesthesia (right) with awake rest in grey. While propofol entails a steepened spectrum as compared to rest, ketamine is associated with spectral flattening.

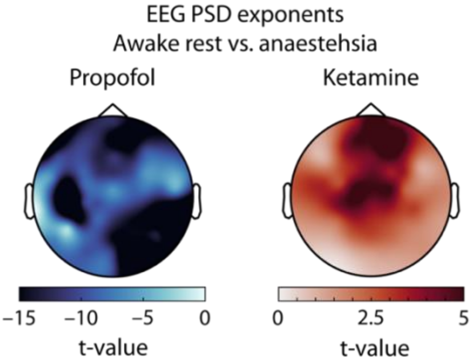


**Figure 1 supplement 2:** Topographically-resolved t-statistics comparing EEG spectral exponents between awake rest and different anaesthetics. Propofol leads to a wide-spread increase in spectral exponents that is present across the entire scalp (left). Ketamine leads to a reduction in spectral exponents that is widely distributed but appears to peak at frontal and central electrodes (right).

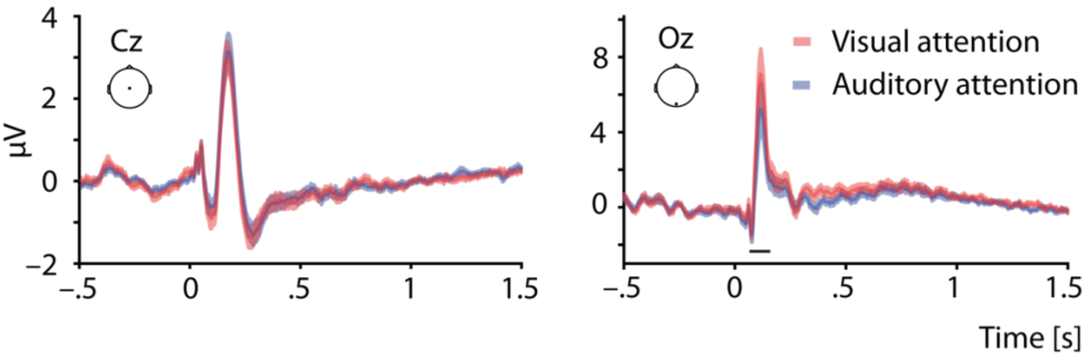


**Figure 3 supplement1:** Evoked responses (ERPs) as a function of attentional focus. Grand average noise onset ERPs (± standard error of the mean, SEM) at electrode Cz (left panel) and electrode Oz (right panel) for visual attention (red) and auditory attention (blue). ERPs are baseline corrected to 500 ms prior to noise onset. Visual attention entailed increased evoked responses at electrode Oz, indicated by the black horizontal line ($p_{FDR} < .05$).

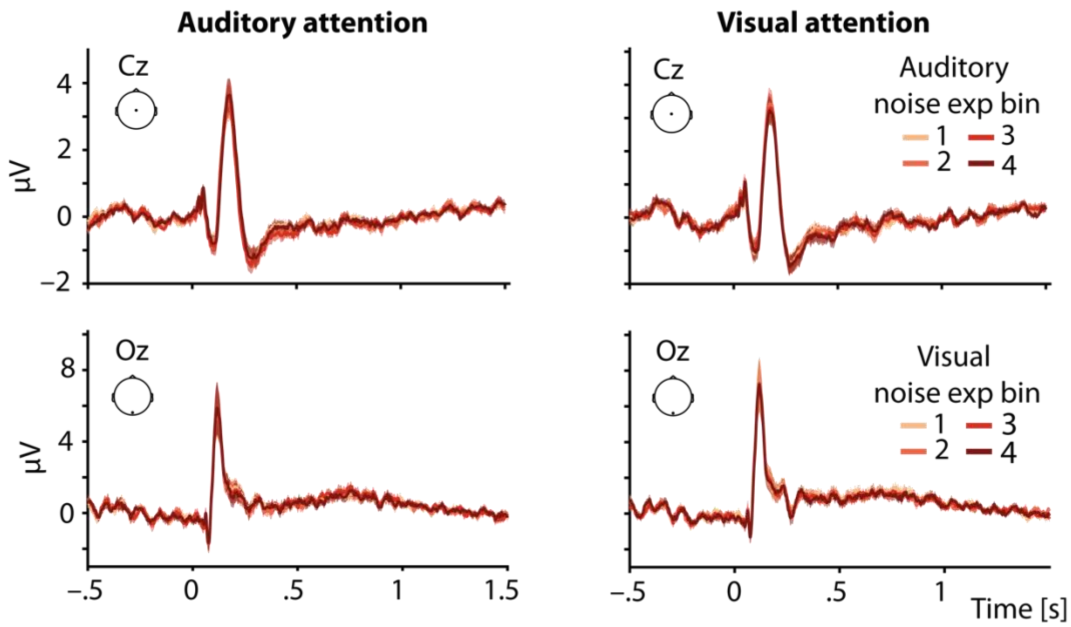


**Figure 4 supplement 1:** Evoked responses (ERPs) as a function of stimulus spectral exponents. (A) Grand average noise onset ERPs (± standard error of the mean, SEM) at electrode Cz (upper panel) and electrode Oz (lower panel) for auditory attention (left column) and visual attention (right column). ERPs are baseline corrected to 500 ms prior to noise onset. Noise onset ERPs (± SEM) for four bins of increasing stimulus spectral exponents (from brighter to darker colours) did not differ significantly.

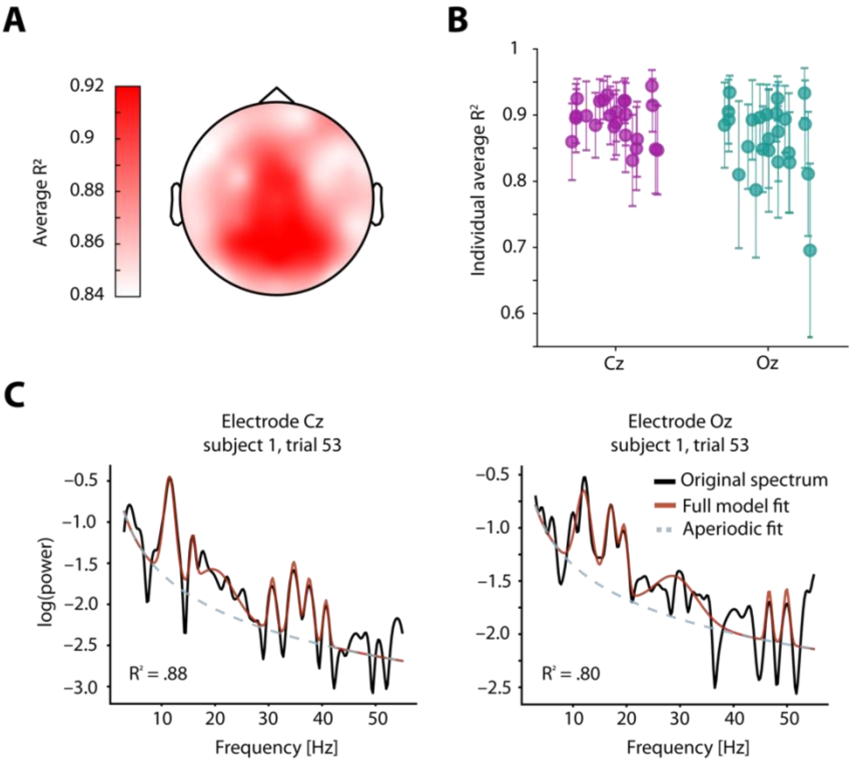


**Figure 4 supplement 2:** Trial-wise spectral parameterization fit statistics and examples. (**A**) Topography of grand average r-squared values from parameterizing single subject EEG spectra. Despite the variation across electrodes, no electrode displayed fits below .84. (**B**) Single subject average (±SD) r-squared values for electrode Cz (left, liac) and electrode Oz (right, teal). (**C**) Representative example of a single trial parameterization result based on data from subject 1, trial 53.

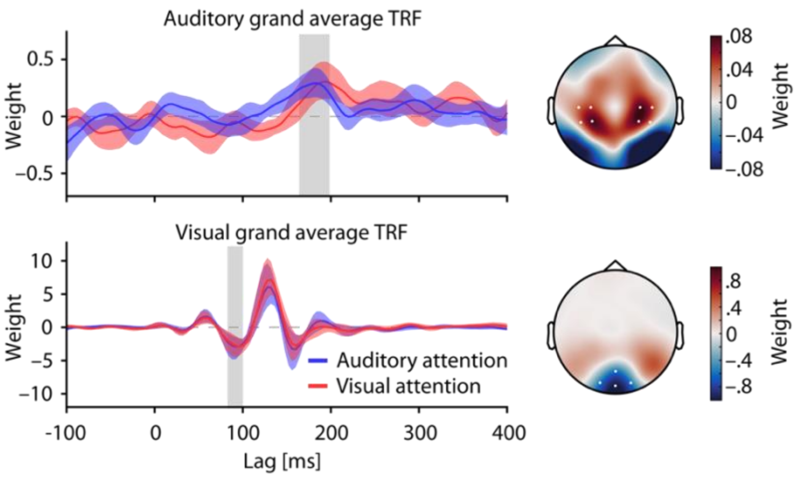


**Figure 4 supplement 3:** Grand average auditory (top) and visual (bottom) temporal response functions for auditory (blue) and visual attention (red). Shaded areas depict bootstrapped 95% confidence intervals (CIs). TRFs were averaged across the electrodes highlighted in the respective topographies. Topographies display weights averaged across time within periods denoted by grey rectangles. Positive lags imply the stimulus preceding the response in time.

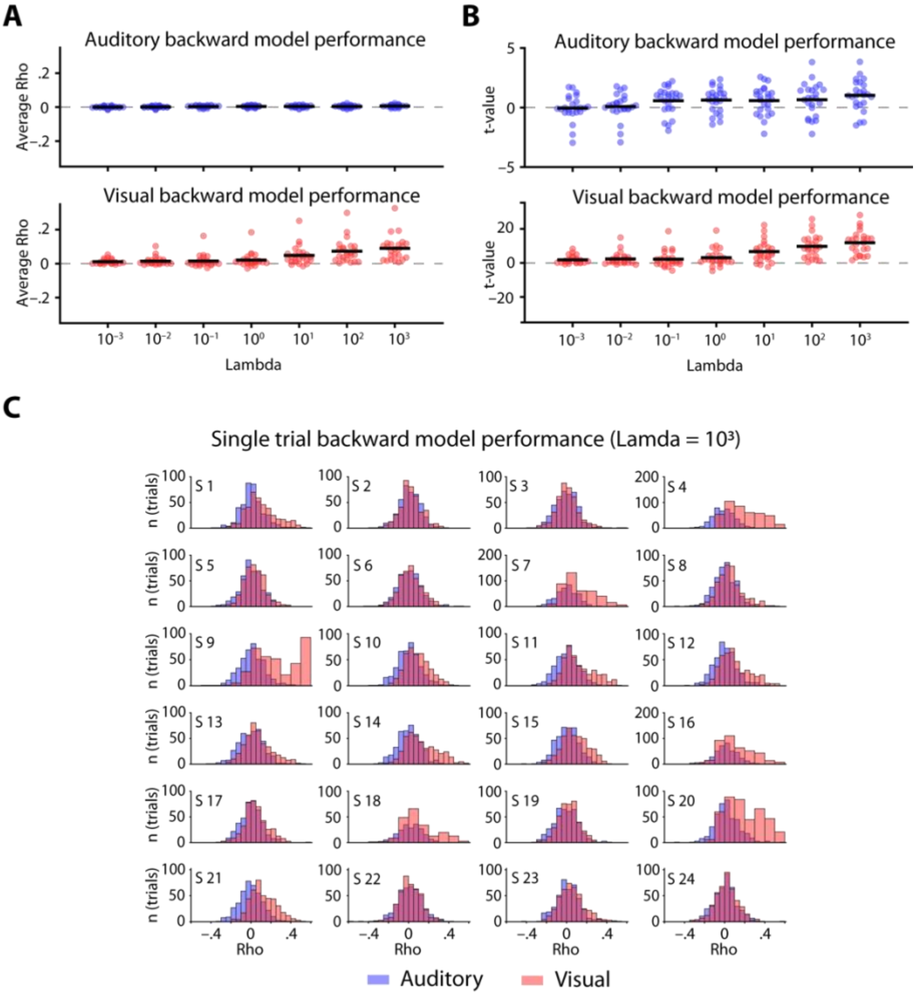


**Figure 4 supplement 4:** Overview of backward model performance. (**a**) Auditory (blue) and visual (red) backward model performance expressed as the average correlation between predicted and observed stimulus time-series across different regularization parameters (Lambda). Dots represent individual Rhos, averaged across trials, horizontal lines denote the mean. (**b**) Individual t-values for auditory and visual backward model performance across different Lambdas, capturing the comparison of individual correlation distributions against zero. Note the difference in y-axes. (**c**) Subject-wise distributions of single trial correlations between predicted and observed stimuli for auditory (blue) and visual (red) stimuli at a Lambda of $10^3$.

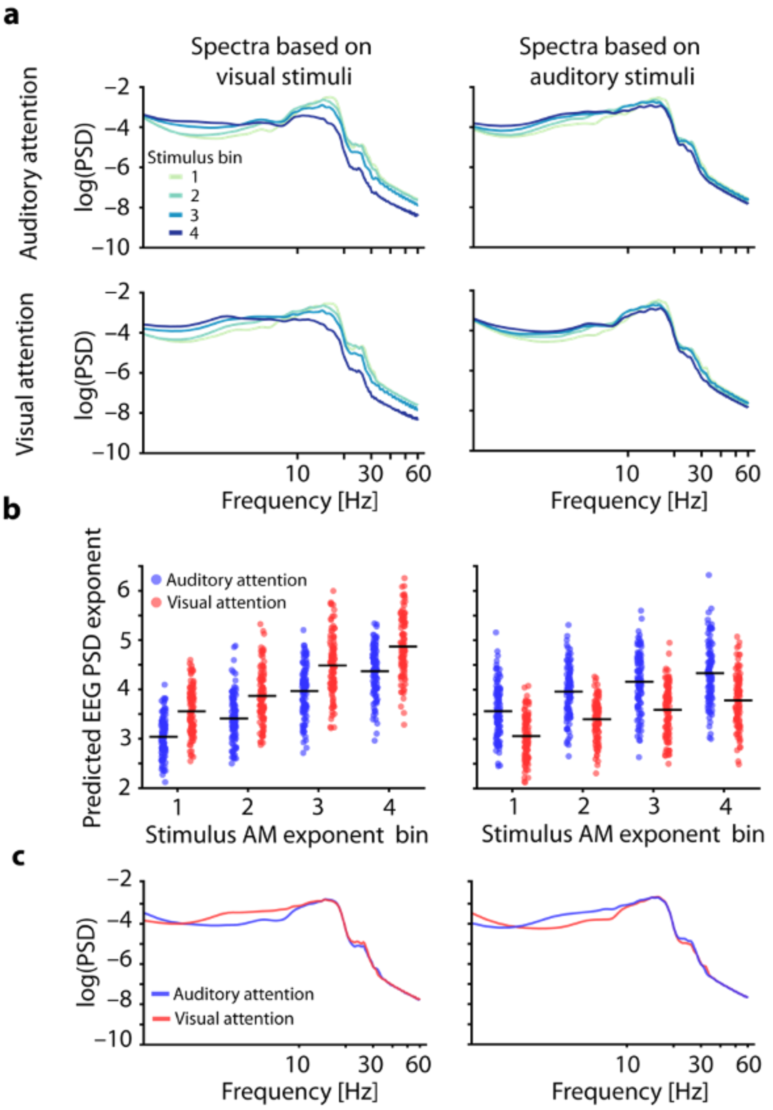


**Figure 4 supplement 5:** Predicted EEG spectra. (**a**) Power spectra of predicted EEG signals based on visual stimuli (left) and auditory stimuli (right). Separate predictions based on canonical TRFs of auditory attention (top) and visual attention (bottom). Spectra are averaged within 4 bins of increasing AM stimulus exponent. (**b**) PSD exponents of predicted spectra based on visual (left) and auditory stimuli (right) and for visual (red) and auditory attention (blue). Dots represent single trial estimates, horizontal lines denote across trial averages. Note the positive link between stimulus and EEG spectral exponent as well as the change of predicted EEG exponents with attentional focus. (**c**) Average power spectra of precited EEG, based on visual (left) and auditory stimuli (right), displayed for visual (red) and auditory attention (blue). Note that a match of predicting stimulus modality and attended modality results in a steepening of spectra.

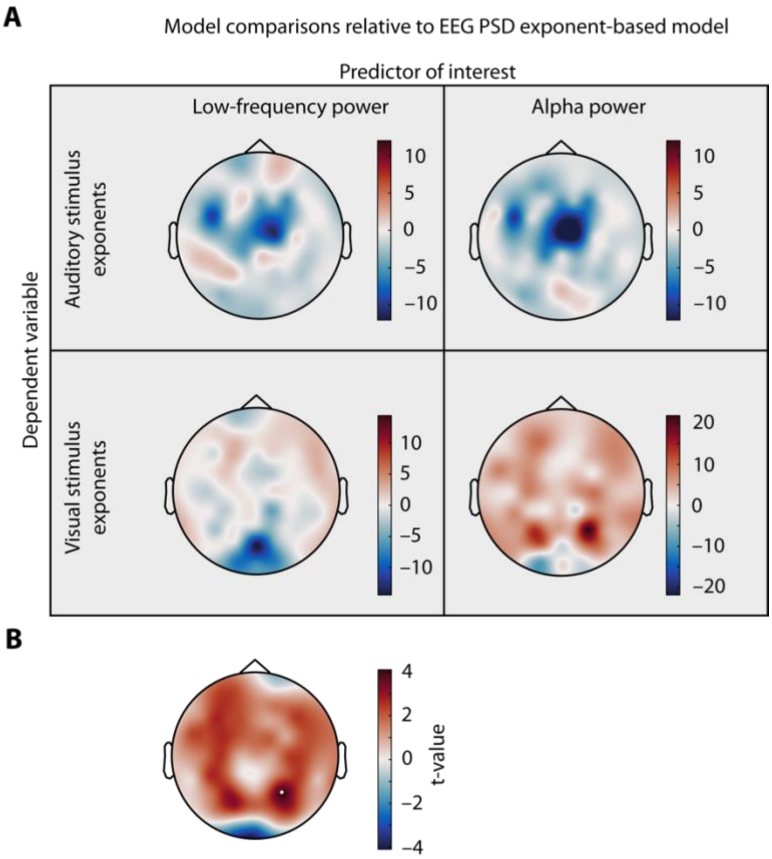


**Figure 4 supplement 6:** Model comparison topographies. (a) Single trial auditory (upper row) or visual stimulus exponents (lower row) were modelled based on electrode-wise low-frequency power (left column) or alpha power (right) column, among other covariates. Models were compared to a model of same size that only differed in the main predictor that consisted of single trial EEG spectral exponents. Topographies display the likelihood ratio test statistic, illustrating no improvements in model fit compared to EEG spectral exponent-based models in all but one model family, illustrating the unique predictive power of aperiodic EEG activity in this context. Alpha power at one parietal electrode explained significantly more variance in visual stimulus exponents. (b) T-values representing the main effect of alpha power on visual stimulus exponents. Highlighted electrode represents p< .05 after FDR correction.

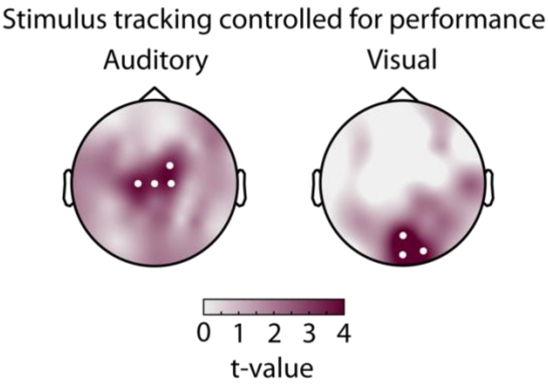


**Figure 4 supplement 7:** Mixed model results, controlling for single trial performance. Topographies depict t-values for the main effect of stimulus spectral exponent, taken from a mixed model of EEG spectral exponents. White dots represent electrodes with significant effects after Bonferroni correction. Auditory stimulus tracking (left) clusters at central electrodes, visual stimulus tracking (right) ad occipital electrodes. Models were identical to tables S1 and S2 but additionally included main effects of single trial accuracy and interactions with auditory and visual stimulus exponents, respectively.

**Table S1: Stimulus tracking model electrode Cz**

| | **EEG spectral exponent** | | | | |
|---|---|---|---|---|---|
| *Predictors* | *Estimates* | *std. Error* | *CI* | *t-value* | *p* |
| Intercept | 1.743 | 0.259 | 1.236 – 2.251 | 6.731 | **<0.001** |
| Auditory spectral exponent | 0.012 | 0.003 | 0.006 – 0.017 | 4.243 | **<0.001** |
| Attention | -0.013 | 0.006 | -0.025 – -0.001 | -2.172 | **0.0299** |
| Visual spectral exponent | -0.001 | 0.003 | -0.007 – 0.004 | -0.418 | 0.6758 |
| Trial number | 0.000 | 0.000 | 0.000 – 0.000 | 4.381 | **<0.001** |
| Resting state EEG exponent | -0.206 | 0.175 | -0.549 – 0.138 | -1.171 | 0.2415 |
| Auditory spectral exponent x Attention | -0.005 | 0.003 | -0.010 – 0.001 | -1.778 | 0.0754 |
| Visual spectral exponent x Attention | -0.003 | 0.003 | -0.008 – 0.003 | -0.948 | 0.3434 |
| **Random Effects** | | | | | |
| $\sigma^2$ | 0.08 | | | | |
| $\tau_{00\ Sub}$ | 0.04 | | | | |
| $\tau_{11\ Sub.Attention}$ | 0.00 | | | | |
| $\rho_{01\ Sub}$ | 0.37 | | | | |
| $N_{Sub}$ | 24 | | | | |
| Observations | 9940 | | | | |
| Marginal $R^2$ / Conditional $R^2$ | 0.020 / 0.332 | | | | |

**Table S2: Stimulus tracking model electrode Oz**

| | EEG spectral exponent | | | | |
|---|---|---|---|---|---|
| *Predictors* | *Estimates* | *std. Error* | *CI* | *t-value* | *p* |
| Intercept | 1.152 | 0.244 | 0.674 – 1.630 | 4.719 | **<0.001** |
| Auditory spectral exponent | 0.006 | 0.003 | -0.000 – 0.012 | 1.865 | 0.0622 |
| Attention | -0.040 | 0.009 | -0.058 – -0.023 | -4.479 | **<0.001** |
| Visual spectral exponent | 0.013 | 0.003 | 0.007 – 0.019 | 4.100 | **<0.001** |
| Trial number | 0.000 | 0.000 | 0.000 – 0.000 | 3.250 | **0.0012** |
| Resting state EEG exponent | 0.001 | 0.186 | -0.363 – 0.365 | 0.005 | 0.9960 |
| Auditory spectral exponent x Attention | 0.003 | 0.003 | -0.003 – 0.009 | 0.927 | 0.3538 |
| Visual spectral exponent x Attention | 0.003 | 0.003 | -0.003 – 0.009 | 1.015 | 0.3100 |
| **Random Effects** | | | | | |
| $\sigma^2$ | 0.10 | | | | |
| $\tau_{00\ Sub}$ | 0.09 | | | | |
| $\tau_{11\ Sub.Attention}$ | 0.00 | | | | |
| $\rho_{01\ Sub}$ | 0.13 | | | | |
| $N_{Sub}$ | 24 | | | | |
| Observations | 9940 | | | | |
| Marginal $R^2$ / Conditional $R^2$ | 0.010 / 0.497 | | | | |
